# Synthesis and biological evaluation of glutathione-responsive 2-alkoxycarbonyl allyl niclosamide prodrugs as anticancer agents

**DOI:** 10.1101/2025.08.20.671294

**Authors:** Anuj K. Singh, Zachary S. Gardner, Greeshma P. Kumpati, Mathew Jacob, Conor T. Ronayne, Venkatram R. Mereddy

## Abstract

Reprogrammed mitochondrial metabolism is recognized as an important target for anticancer therapy. Niclosamide, an FDA approved anthelmintic agent with mitochondrial uncoupling activity, has shown promise as an anticancer agent. However, off target mitochondrial toxicity has rendered the utility of this agent at clinically effective doses. Here, we synthesize a variety of prodrugs on the niclosamide template based on the Baylis-Hillman (BH) reaction, with the hypothesis that niclosamide will be released upon the reaction with cellular nucleophiles, including thiols. Consistent with this hypothesis, the BH-prodrugs release the parent niclosamide in the presence of cysteine and glutathione, and the cancer cell proliferation inhibition properties of the lead candidates are retained when compared to the parent niclosamide. Mitochondrial respiration assays illustrate that niclosamide acutely uncouples the mitochondria, and the lead BH-prodrug **6b** does not, providing evidence of this prodrug strategy in mitigating off target toxicities. In a dose escalation study, the lead candidate **6b** is generally well tolerated in healthy mice as evidenced by zero mortality, normal grooming pattern, and normal weight gains. Finally, **6b** exhibits 54% volume tumor growth inhibition properties in a syngraft model of breast cancer in mice. The studies herein provide a novel methodology for the application of BH prodrugs on the niclosamide template and the anticancer applications.

## INTRODUCTION

Altered cellular metabolism is a well-recognized hallmark of cancer, enabling rapid proliferation through reprogrammed energy production, and biosynthetic pathways **[1–7].** Niclosamide, a salicylanilide-based FDA-approved anthelmintic agent, has attracted considerable attention due to its ability to target multiple oncogenic signalling pathways, including NF-κB **[14]**, Wnt/β-catenin **[15]**, and reactive oxygen species (ROS) modulation **[16]**. These molecular effects provide a compelling rationale for repositioning niclosamide as a potential anticancer agent **[17–20].** In this context, drug repurposing has emerged as a powerful strategy to identify new anticancer agents from previously approved drugs with known safety profiles **[8–13].**

Recent studies have revealed that niclosamide functions as a potent mitochondrial uncoupling agent **[21–24].** By dissipating the proton gradient across the inner mitochondrial membrane, niclosamide disrupts ATP synthesis via oxidative phosphorylation **[25–27].** This uncoupling impairs mitochondrial bioenergetics, elevates ROS production, and can activate intrinsic apoptotic pathways **[23, 24].** These mitochondrial effects have been shown to selectively impair cancer cell viability, especially in metabolically stressed or oxidative phosphorylation-reliant tumors **[22, 24].** The therapeutic potential of niclosamide as a metabolic disruptor positions it as a promising candidate for targeting mitochondrial vulnerabilities in cancer cells **[24–26].**

Despite its broad pharmacological activity and high tolerability in both animals and humans, niclosamide’s clinical translation for cancer therapy is limited by poor oral bioavailability, extensive first-pass metabolism, and rapid systemic efflux **[17, 27].** These challenges significantly reduce its effective plasma concentrations and therapeutic index **[26, 27]**. One strategy to overcome these pharmacokinetic limitations is to design prodrugs that block the phenolic OH group of niclosamide **[27]**. Masking this key functional group can enhance lipophilicity, may improve metabolic stability, thereby improving systemic exposure and therapeutic potential **[28]**.

The Baylis–Hillman (BH) reaction is an important C–C bond-forming transformation that generates functionalized allyl alcohols and amines under mild conditions **[28–31]**. Owing to its operational simplicity, atom economy, and the versatility of BH-derived synthons, this reaction has found applications in medicinal chemistry **[28, 29]**. The acetates of BH alcohols undergo facile allylic rearrangement via an SN2′ mechanism with various nucleophiles— including carbon, sulphur, nitrogen, and oxygen to yield tri-substituted α,β-unsaturated systems **[28, 32]**. Our interest in exploring BH chemistry as a scaffold for novel anticancer agents led us to integrate this platform with niclosamide. In previous reports, bromomethyl acrylates derived from BH alcohols exhibited reactivity toward intracellular nucleophiles and exhibited significant anticancer efficacy **[28, 32]**. In this work, we designed Baylis–Hillman- based niclosamide prodrugs to address some of its limitations.

The rationale behind this approach was to exploit the mitochondrial localization and redox sensitivity of cancer cells by developing glutathione-responsive BH-based prodrugs **[28, 32]**. Notably, cancer cells maintain significantly higher intracellular concentrations of glutathione (GSH) compared to normal cells, a feature that enables them to neutralize oxidative stress and maintain redox homeostasis **[33]**. This elevated GSH level, while protective, also presents a selective vulnerability that can be harnessed by redox-sensitive therapeutic agents **[33]**. By linking niclosamide with BH acrylates, the resulting prodrugs were designed to undergo nucleophilic cleavage in this reductive intracellular environment, particularly within tumor cells, thereby releasing the active drug in a site-selective manner. The mitochondrial release of niclosamide was anticipated to cause proton gradient collapse, bioenergetic failure, and disruption of biosynthetic processes, ultimately impeding tumor growth. This prodrug strategy was aimed at enhancing the tumor-selective activation of niclosamide’s anticancer effects **[28–32]**.

## RESULTS AND DISCUSSION

### Design and synthesis of niclosamide prodrugs

Niclosamide, a phenolic protonophore, perturbs a wide variety of cellular and molecular processes stemming from its mitochondrial uncoupling characteristics. To limit off-target effects of niclosamide, we sought to design a prodrug that renders the uncoupling capacity of niclosamide inactive until reacted with intratumoral nucleophiles including glutathione. To facilitate this prodrug design strategy, we explored the development of BH reaction-derived prodrugs on the niclosamide template. We have shown the utility of BH chemistry with simple substrates including phenols, carboxylic acids, and amines exhibiting release of the parent compounds upon interaction with nucleophilic cellular components through mechanisms such as SN2 or SN2’ reactions, and possibly 1,4-addition reaction (**Scheme 1**) **[28]**. Here, we leveraged the nucleophilic substitution reactivity of niclosamide to design prodrugs of bromomethyl acrylate **4a-4d** (**Scheme 1A**). These acrylates can be readily obtained via BH reaction of formaldehyde with acrylates followed by bromination of the resulting alcohols (**Scheme 1A**). Specifically, niclosamide phenolic group served as the nucleophile, facilitating the displacement of bromine from methyl 2-(bromomethyl)acrylate (**Scheme 1B**). This reaction resulted in the formation of methyl 2-((4-chloro-2-((2-chloro-4-nitrophenyl) carbamoyl) phenoxy) methyl) acrylate **6a**. Similarly, other alkyl bromides were reacted with niclosamide in a similar fashion, enabling the synthesis of additional BH reaction derived prodrugs **6b**, **6c** and **6d**. These prodrugs hold potential for selective release of niclosamide upon reaction with nucleophilic cellular components such as glutathione (**Scheme 1C**). In general, the concentration of glutathione is several folds higher in cancer cells than plasma concentration **[28]** and this fact may allow the selective release of niclosamide in cancer cells.

**Scheme 1.**
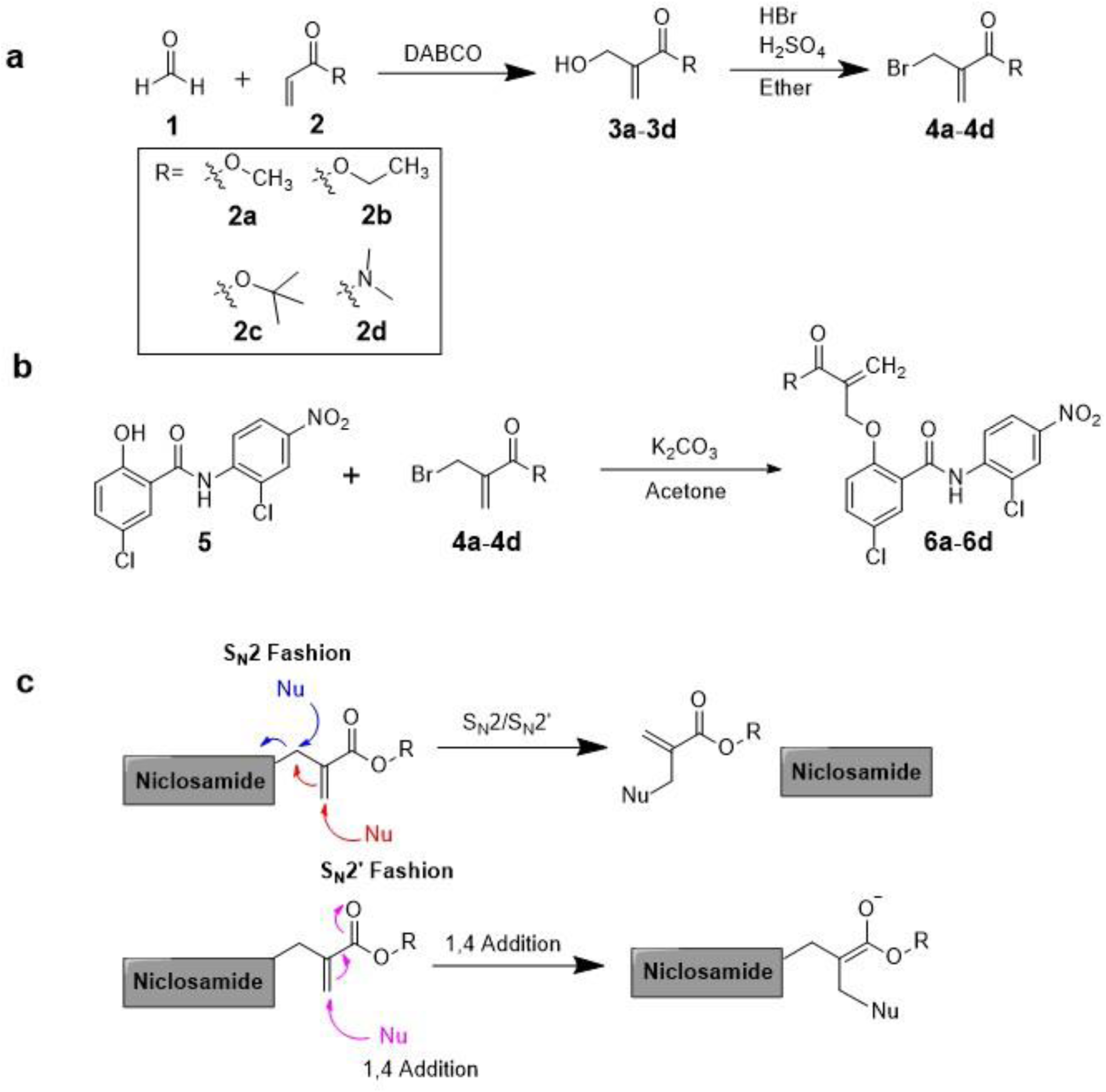
(**A**) Baylis-Hillman reaction derived alcohols and corresponding bromides for the synthesis of (**B**) niclosamide prodrugs 6a-6dm via standard nucleophilic substitution onto the phenol of niclosamide 5. (**C**) Reaction scheme of BH-derived niclosamide prodrugs that can form adducts with intracellular nucleophiles in an S_N_2, S_N_2’, and or 1,4-addition fashion.

### Baylis-Hillman-derived niclosamide prodrugs **6a-6d** inhibit cancer cell proliferation

To evaluate the anticancer activity of niclosamide **5** and prodrugs **6a-6d**, we assessed the compounds effects on breast cancer cell viability *in vitro*. We used a panel of breast cancer cells including murine isogeneic 67NR and 4T1 lines, along with human triple negative breast cancer MDA-MB-231. Here, using the 3-(4,5-di methyl thiazol-2-yl)-2,5-diphenyltetrazolium bromide (MTT) assay, we determined that niclosamide **5** inhibited cancer cell proliferation with IC_50_ values ranging from 0.23 – 5.10 µM across the cell lines tested (**Table 1**). Interestingly, the methyl **6a** and ethyl **6b** BH-ester pro drugs had modest increases potency (0.19 – 2.76 µM), and the tert-butyl ester **6c** and amide **6d** exhibited modest decreases in activity across the cell lines tested (0.64 - >100 µM, **Table 1**). These results illustrate, overall, that the potency of the BH-prodrugs are largely conserved when compared to the parent niclosamide **5**, and the resulting potency may be a reflection of the BH-prodrug releasing the active drug. It can be noted that all of the tested compounds had exaggerated effects on 4T1 when compared to 67NR and MDA-MB-231 cells, which may highlight the dependence of 4T1 on mitochondrial respiration for maintaining cell viability in the presence of mitochondrial uncoupling agents like niclosamide.

**Table 1.**
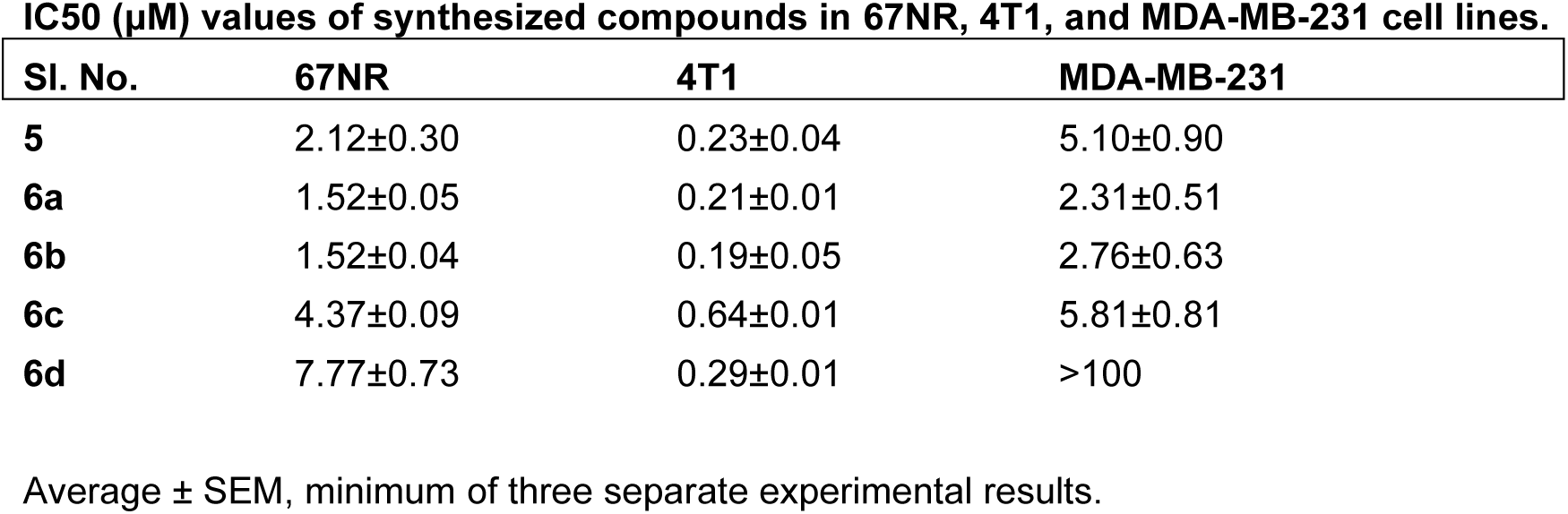
IC_50_ values of synthesized compounds in 67NR, 4T1, and MDA-MB-231 cell lines. Data represents the average ± SEM of three independent experiments (n=3).

| Sl. No. | 67NR | 4T1 | MDA-MB-231 |
| --- | --- | --- | --- |
| **5** | 2.12±0.30 | 0.23±0.04 | 5.10±0.90 |
| **6a** | 1.52±0.05 | 0.21±0.01 | 2.31±0.51 |
| **6b** | 1.52±0.04 | 0.19±0.05 | 2.76±0.63 |
| **6c** | 4.37±0.09 | 0.64±0.01 | 5.81±0.81 |
| **6d** | 7.77±0.73 | 0.29±0.01 | >100 |

Average ± SEM, minimum of three separate experimental results.

### Lead candidate **6b** releases niclosamide in the presence of glutathione and cysteine

The candidate prodrugs have been designed to react with intracellular nucleophiles, particularly thiol residues, in an SN2/SN2’ fashion. This reaction will result in the parent niclosamide **5** being substituted and released as a leaving group. To test this hypothesis, we used thin-layer chromatography to examine the reactivity of one of the lead candidates **6b** with thiols cysteine and glutathione. In this experiment, the appearance and ratio of prodrug **6b** to parent niclosamide **5** can be monitored after incubation with a nucleophile. Separate spots under UV were observed as the polarity of **6b** and **5** are distinct and resolve to distinguishable relative fronts (Rf) on the TLC silica plate (**Figure 1**). These results indicate that the prodrug **6b** reacted with thiols cysteine and glutathione, as parent niclosamide **5** was released which was evidenced by the appearance of parent niclosamide **5** UV spot under these conditions (**Figure 1**). Additionally, these results illustrate the importance of the thiol in facilitating the prodrug release as alanine and serine amino acids were not sufficient to promote the release of niclosamide **5** (**Figure 1**).

**Figure 1.**
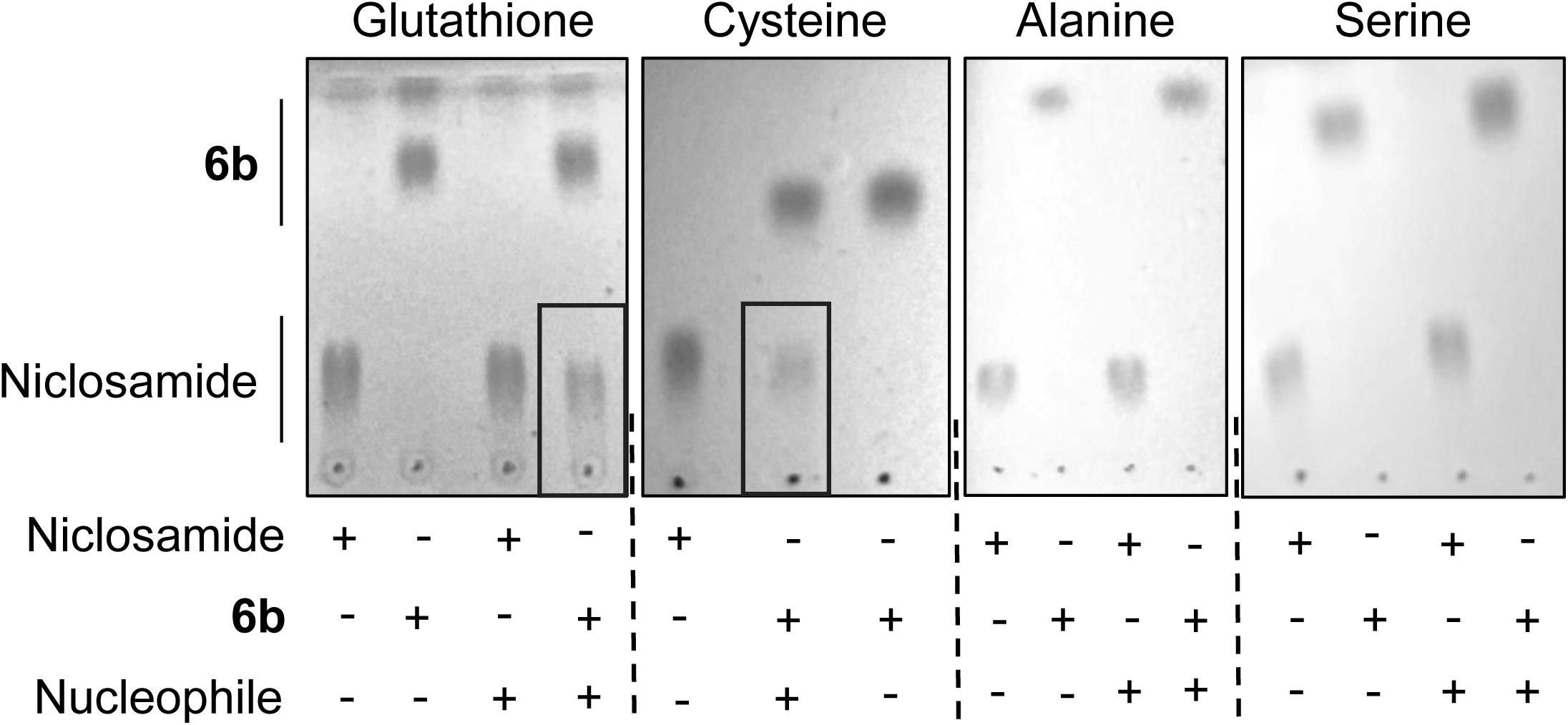
Thin-layer chromatography experiments illustrate that prodrug **6b** releases parent niclosamide **5** in the presence of thiol-nucleophiles glutathione and cysteine, but not in the presence of non-nucleophilic alanine, or oxygen-nucleophile serine. Note the boxed re-appearance of niclosamide spot in the presence of glutathione and cysteine. Images are representative of 2 or more experiments for each condition (n>2).

### BH-Prodrug **6b** exhibits attenuated effects on mitochondrial respiration as per the parent niclosamide **5**

The rationale for synthesizing a prodrug of niclosamide that blocks the phenol group is to temporarily inhibit the uncoupling activity of these drugs until the parent niclosamide is released (**Scheme 1**). In this regard, we sought to test this hypothesis by evaluating the effects of prodrug **6b** on the mitochondrial respiration when compared to the known uncoupling activity of the parent niclosamide. Here, we performed a mitochondrial stress test in 67NR breast cancer cells to evaluate acute mitochondrial perturbations of **6b** compared to niclosamide **5** (**Figure 2**). In this experiment, oxygen consumption rates were first measured to determine basal respiration rates, which were equal across test conditions (**Figure 2A-B**). Test compounds (**6b** or niclosamide **5**) were then injected at various concentrations and acute effects on mitochondrial respiration were assessed. Injection of niclosamide **5** resulted in an acute increase in respiration at intermediate concentrations, consistent with its ability to uncouple the mitochondrial respiration in a mitohormetic fashion (**Figure 2A, C**). As expected, blocking the phenolic group with BH-prodrug in **6b** attenuated the ability of **6b** to uncouple the mitochondria, as respiration in 67NR cells treated with **6b** was increased to a lesser extent and at higher concentration ranges when compared to niclosamide **5** (**Figure 2A, C**). Cells were then exposed to ATP synthase inhibitor oligomycin, where the extent of decreases in OCR can be attributed to a complementary balance between proton leak across the inner membrane, and oxygen consumed for ATP production. It was observed that niclosamide potently inhibited ATP production and led to a corresponding proton leak (**Figure2A, D-E**). In comparison, **6b** did not drastically affect ATP production, with only modest but statistically significant decreases at only 1 µM (**Figure 2A, D**). Interestingly, **6b** induced proton leak at all concentrations tested, which may be a reflection of its ability to uncouple the mitochondria at these concentrations, albeit to a lesser extent than niclosamide **5** (**Figure 2A, E**). Injection with mitochondrial uncoupling agent FCCP then prompted cells to work at maximal OCR, and it was noted that niclosamide **5**, but not **6b** affected maximal and/or spare respiratory capacities (**Figure 2A, F-G)**. Respiration was finally eliminated with the combined injection of rotenone and antimycin A to inhibit complex I and III, respectively. These results clearly illustrate that protecting the phenol of niclosamide with a BH prodrug attenuates the uncoupling activity of this drug, consistent with our design hypothesis to mitigate off-target toxicities associated with non-targeted uncoupling activity. The shift in both concentration ranges and magnitude of the effects of **6b** on the mitochondria when compared to niclosamide **5** likely represent a small portion of the prodrug **6b** reacting to release niclosamide within the time-scale of this experiment. Combined with the data in **Figure 1** that illustrates the ability of prodrug **6b** to release niclosamide under conditions of glutathione or cysteine nucleophiles, qualifies this candidate for further investigation as an anticancer agent in mouse models of breast cancer.

**Figure 2.**
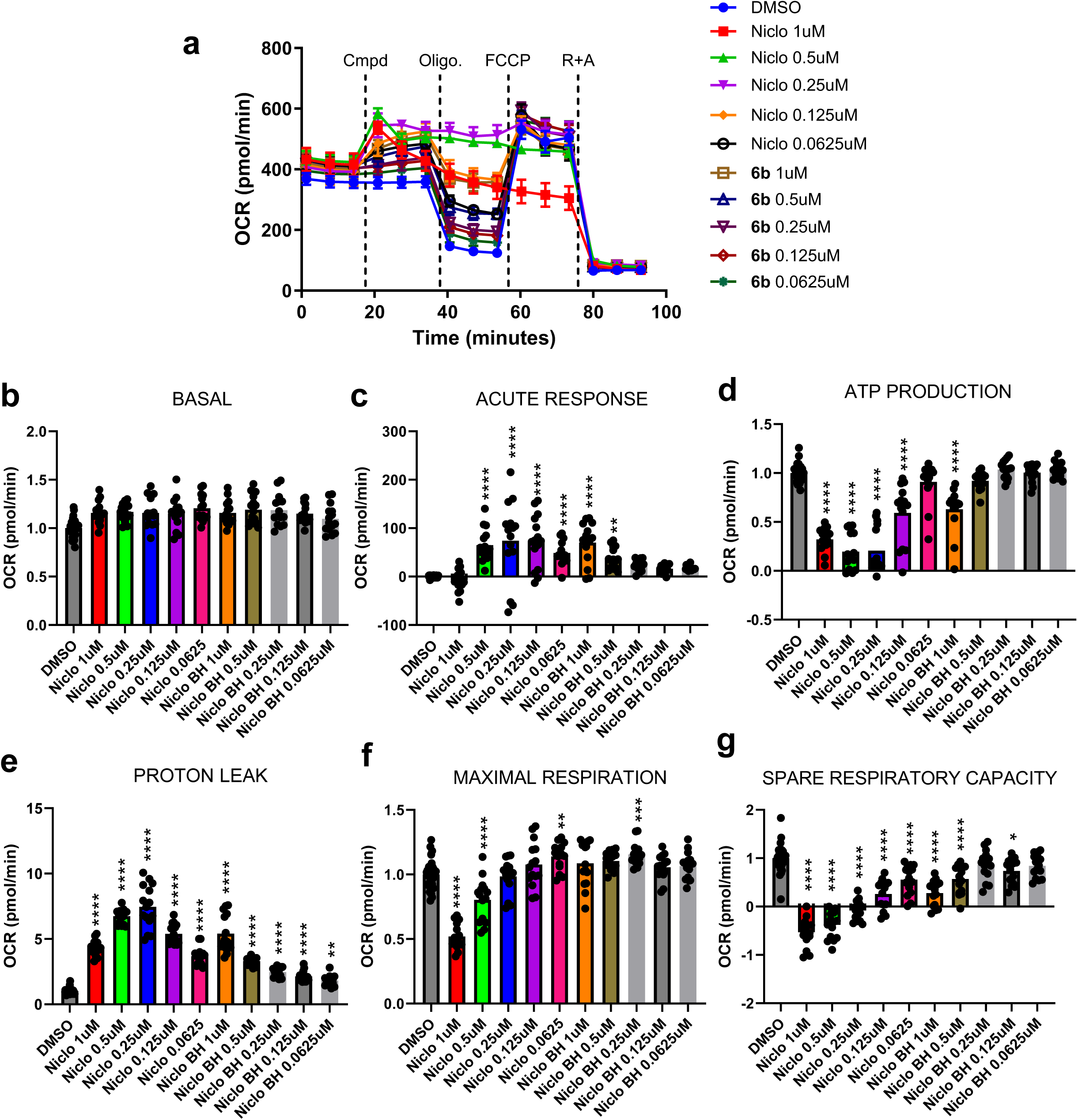
Seahorse XFe assays illustrate attenuated mitochondrial effects of **6b** when compared to niclosamide **5**. (**A**) Mitochondrial respiration profile and corresponding oxygen consumption rates (OCR) of 67NR cells subsequently treated with indicated test compounds (Cmpd), oligomycin (Oligo. 1 µM), FCCP (250 nM), and a cocktail of rotenone and antimycin A (R + A, 0.5 µM respectively). Changes in OCR in response to specific injections under each test compound condition were used to calculate parameters in B-G. (**B**) Basal respiration calculated as respiration of cells in each condition prior to compound injection. (**C**) Acute response calculated as OCR induced after Cmpd injection when compared to basal. (**D**) ATP production calculated as OCR change after oligomycin injection. (**E**) Proton leak calculated as the complementary difference between basal and ATP production rates. (**F**) Maximal respiration calculated as the difference between post-oligomycin respiration and post FCCP-stimulated respiration. (**G**) Spare respiratory capacity as calculated by the difference between basal and maximal rates. Note the shift in mitohormetic response of niclosamide **5** at intermediate concentrations, to high concentrations in **6b** treated cultures. Data is pooled from three independent experiments (N=3), representing 30 and 15 technical replicates for DMSO, and compound treated conditions, respectively (n=30, 15). Statistics were calculated using one-way ANOVA, where *p<0.05, **p<0.01, ***p<0.001, and ****p<0.0001.

### Lead candidate **6b** exhibits tumor-growth inhibition properties comparable to that of niclosamide

To evaluate the ability of lead candidate **6b** to inhibit tumor growth *in vivo,* we tested the tumor growth inhibition properties of **6b** in a syngraft model of breast cancer 67NR in BALB/c mice. First, we determined the maximum tolerated dose (MTD) of our candidate **6b** and niclosamide **5** where mice tolerated both compounds at equal doses when escalated from 2.5mg/kg to 30mg/kg (**Figure 3A**). Then, using the pre-determined MTD, we performed the 67NR syngraft model. Here, we inoculated healthy female BALB/c mice with 67NR cells into the right flank and tumors were allowed to reach ∼100mm^3^. Mice were then randomly separated into three groups (n=8); 1) vehicle (10%DMSO/saline), 2) lead candidate **6b** (10 mg/kg, once daily, i.p.), and 3) niclosamide **5** (10 mg/kg, once daily, i.p.). Mice were treated with drug candidates for 9 days, after which the aggressive nature of the vehicle-treated tumors qualified for study termination. Tumor volumes were recorded over the duration of the study where it was determined that BH-prodrug **6b** inhibited tumor volumes by 54%, when compared to 19% of parent niclosamide, albeit these volumes were found to be statistically insignificant (**Figure 3B&C**). After the study, tumors were resected and tumor masses were obtained where **5** and **6b** exhibited statistically significant tumor mass reductions of 49% and 50% when compared to vehicle, respectively (**Figure 3D**). This preliminary *in vivo* tumor growth inhibition study provides evidence that the BH-prodrug strategy of **6b** retained activity when compared to the parent niclosamide.

**Figure 3.**
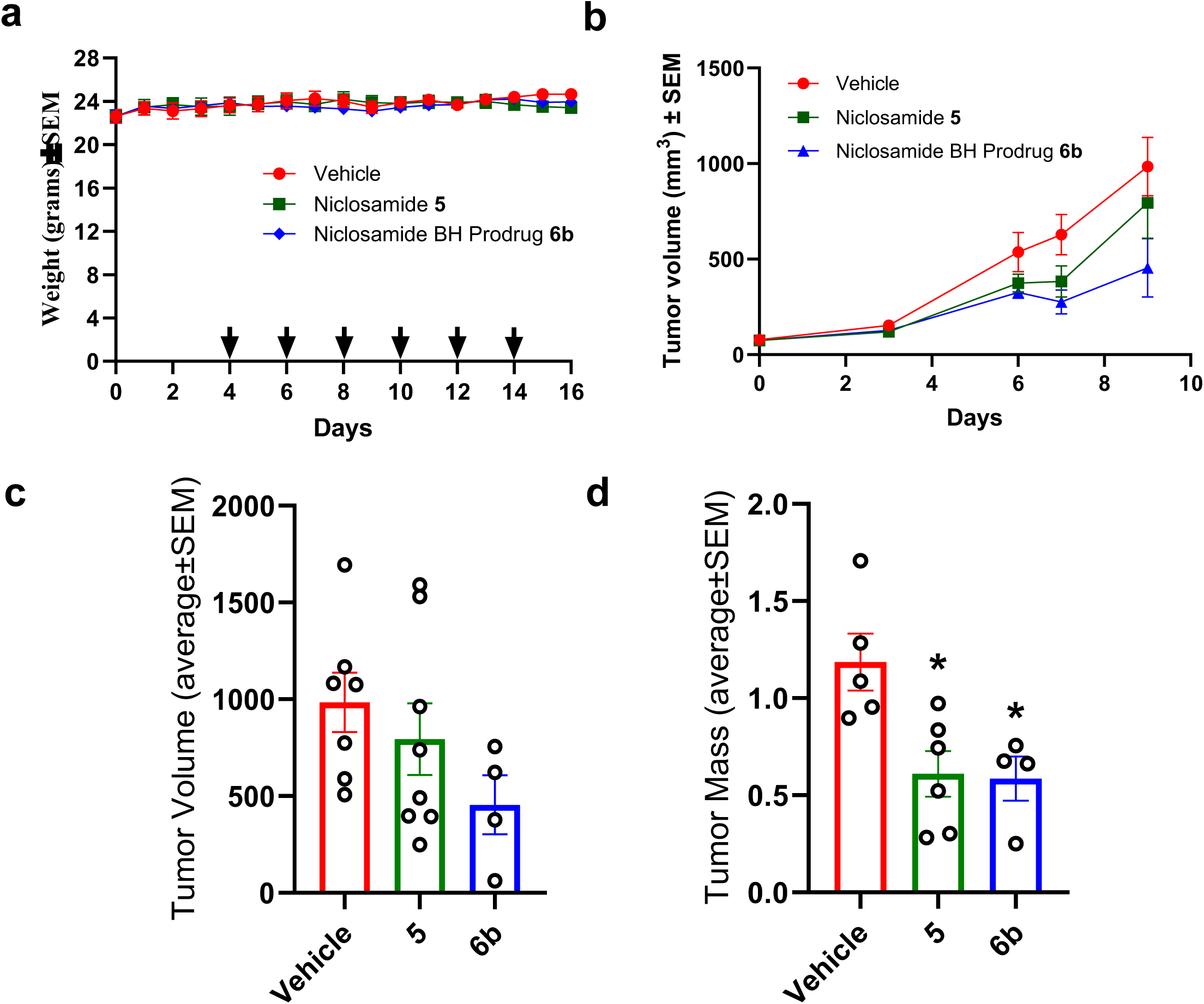
Prodrug **6b** exhibits tumor growth inhibition properties in a 67NR model of breast cancer in mice. (**A**) Maximum tolerated dose of compounds **5** and **6b** in healthy CD-1 mice (n=5) up to 30 mg/kg. Mice were initiated at a 2.5 mg/kg dose, which was incrementally increased to 5, 10, 15, 20, 25, and finally 30 mg/kg at the days indicated with arrows. (**B**) 67NR model in BALB/c mice (n=8) indicates trending increased mean tumor volumes in mice treated with **6b** (10mg/kg, i.p. once daily) when compared to **5** (10mg/kg, i.p. once daily) and vehicle. Data represents the average ± SEM of end-point tumor volumes (**C**) and endpoint tumor masses (**D**). Statistical significance was calculated using the Mann-Whitney test (*p<0.05).

## CONCLUSIONS

In the current study, we synthesized a few prodrugs on the niclosamide based on the BH reaction derived bromomethyl acrylates and amides. Cancer cell proliferation inhibition properties of the lead candidates are retained when compared to the parent niclosamide against a series of breast cancer cell lines *in vitro*. Consistent with the hypothesis, the BH- prodrugs released the parent niclosamide in the presence of thiol nucleophiles such as cysteine and glutathione. Mitochondrial respiration assays illustrated that niclosamide acutely uncoupled the mitochondria, and the lead BH-prodrug **6b** did not, providing evidence of this prodrug strategy in mitigating off target toxicities. Finally, the dose escalation studies and anticancer efficacy studies of the lead candidate **6b** was well tolerated and exhibited 54% tumor volume growth inhibition properties in a syngraft model of breast cancer in mice. The studies herein offer a novel application of BH derived bromomethyl acrylates on the niclosamide template and a new methodology for anticancer applications.

## METHODS

### Materials

Niclosamide (5-chloro-N-(2-chloro-4-nitrophenyl)-2-hydroxybenzamide) (Sigma-Aldrich), formaldehyde (Sigma-Aldrich), DABCO (Sigma-Aldrich), methyl acrylate (Sigma-Aldrich), ethyl acrylate (Sigma-Aldrich), tert-butyl acrylate (Sigma-Aldrich), N, N-dimethylacrylamide (Sigma-Aldrich). Other chemical reagents were of high-grade quality and purchased from Sigma-Aldrich (St. Louis, MO).

The 1H- and 13C-NMR spectra were plotted on a Varian Oxford-500 and Bruker AscendTM 400 spectrometers. High resolution mass spectra (HRMS) were recorded with a Bruker micrOTOP-Q III ESI mass spectrometer.

### Representative procedure for the synthesis of 2-(bromomethyl)acrylates and acrylamide derivatives

To an aldehyde 1 (2mmol) was added an activated alkene acrylates 2a-2d (2.4mmol) and stirred in the presence of DABCO (2mmol) at room temperature. Upon completion (TLC), water was added, and product was extracted with ethyl acetate. Organic layer washed with saturated sodium bicarbonate to remove unreacted components. Organic layer was then dried using MgSO4 and filtered. Crude product was then concentrated under vacuum to obtain corresponding Baylis-Hillman alcohol 3a-3d as a crude product.

The crude BH alcohol 3a-3d was then dissolved in anhydrous ether and treated with hydrogen bromide gas in the presence of catalytic sulfuric acid at 0°C. The reaction was stirred for 2–3 hours, allowing conversion to the bromomethyl derivative. The reaction mixture was quenched with water and extracted with ethyl acetate. The organic phase was washed, dried, and concentrated under reduced pressure. The resulting acrylate was purified by column chromatography to obtain pure compounds 4a-4d.

### Synthesis of methyl 2-((4-chloro-2-((2-chloro-4 nitrophenyl) carbamoyl) phenoxy) methyl) acrylate

To compound 4a (3mmol) in acetone (10mL) was added Niclosamide 5 (2mmol) in the presence of the potassium carbonate (3mmol). Reaction was stirred for 3 hours and confirmed with TLC for the reaction completion. The reaction was then washed with brine solution and ethyl acetate (3x25mL). The organic phase evaporated using a rotatory evaporator and dried under vacuum. This crude was washed with diethyl ether and was vacuum filtered to obtain the pure compound 6a (84%).

### Synthesis of ethyl 2-((4-chloro-2-((2-chloro-4-nitrophenyl) carbamoyl) phenoxy) methyl) acrylate

To compound 4b (3mmol) in acetone (10mL) was added Niclosamide 5 (2mmol) in the presence of the potassium carbonate (3mmol). Reaction was stirred for 3 hours and confirmed with TLC for the reaction completion. The reaction was then washed with brine solution and ethyl acetate (3x25mL). The organic phase evaporated using a rotatory evaporator and dried under vacuum. This crude was washed with diethyl ether and was vacuum filtered to obtain the pure compound 6b (88%).

### Synthesis of tert-butyl 2-((4-chloro-2-((2-chloro-4-nitrophenyl) carbamoyl) phenoxy) methyl) acrylate

To compound 4c (3mmol) in acetone (15mL) was added Niclosamide 5 (2mmol) in the presence of the potassium carbonate (3mmol). Reaction was stirred for 5 hours and confirmed with TLC for the reaction completion. The reaction was then washed with brine solution and ethyl acetate (3x25mL). The organic phase evaporated using a rotatory evaporator and dried under vacuum. This crude was washed with diethyl ether and was vacuum filtered to obtain the pure compound 6c (77%).

### Synthesis of 5-chloro-N-(2-chloro-4-nitrophenyl)-2-((2-(dimethylcarbamoyl) allyl) oxy) benzamide

To compound 4d (3mmol) in acetone (15mL) was added Niclosamide 5 (2mmol) in the presence of the potassium carbonate (3mmol). Reaction was stirred for 3 hours and confirmed with TLC for the reaction completion. The reaction was then washed with brine solution and ethyl acetate (3x25mL). The organic phase evaporated using a rotatory evaporator and dried under vacuum. This crude was washed with diethyl ether, vacuum filtered and purified via column chromatography (1:3, EtOAc:Hexanes) to obtain the pure compound 6d (77%).

### Cell Culture

MDA-MB-231 cells (ATCC) were grown in DMEM supplemented with 10% FBS and penicillin-streptomycin (50U/ml-50µg/ml). MIAPaCa-2 cells (ATCC) were cultured in DMEM supplemented with 10% FBS, 2.5% Horse Serum and penicillin- streptomycin (50U/ml- 50µg/ml). 4T1 cells (ATCC) were cultured in RPMI-1640 supplemented with 10% FBS and penicillin-streptomycin (50U/ml- 50µg/ml). 67NR cells (ATCC) were cultured in RPMI-1640 supplemented with 10% FBS and penicillin-streptomycin (50U/ml-50µg/ml). BxPC-3 cells (ATCC) were cultured in RPMI-1640 supplemented with 10% FBS and penicillin-streptomycin (50U/ml- 50µg/ml).

### Cell Proliferation Inhibition Assay (MTT)

Cells were plated at a density of 5x10^3^ cells per well in 96-well plates and allowed to adhere overnight at 37°C in a 5% CO_2_ atmosphere. The cells were then treated in duplicate with serial dilutions of the test compounds and incubated for an additional 72 hours under the same conditions. Following the treatment period, 0.5 mg/mL of 3-(4,5-dimethylthiazol-2-yl)-2,5- diphenyltetrazolium bromide (MTT) was added to each well and incubated for four hours. The resultant formazan crystals were dissolved using a solution of 10% SDS in 0.01N HCl and further incubated for four hours. Absorbance at 570 nm was measured for each well. The IC_50_ values, representing the concentrations at which cell growth was inhibited by 50%, were determined using untreated wells as the reference for 100% cell survival.

### Thin Layer Chromatography based analysis of Niclosamide and BH Prodrug Conjugation with GSH

Niclosamide and its Baylis–Hillman (BH) prodrug 6b (1 mM each) was incubated with glutathione (GSH), serine, and alanine (10 mM each) in PBS buffer (pH 7.4) at 37 °C for 24 h. Post incubation, 100 µL of ethyl acetate was added to each reaction mixture, followed by extraction. The organic (ethyl acetate) layer was collected and spotted on silica gel TLC plates. Plates were developed using a hexane: ethyl acetate mobile phase and visualized under UV light. Control samples with individual compounds were included to assess specificity of reactivity.

### Seahorse XFe96® Assessment of Glycolysis and Mitochondrial Respiration

Cells were plated at a concentration of 20,000 cells/well in a Seahorse XFe96 culture microplate (Agilent, part no. 101085-004) and incubated for 24 hours at 37°C at 5% CO_2_. Sensor cartridge (Agilent, part no. 102416-100) was hydrated with XF calibrant (Agilent, part no. 100840-000) overnight at 37°C in an incubator without CO_2_. The assay media was prepared from Seahorse base medium (Agilent, part no. 102353-100) and enriched with glucose (10mM), glutamine (1mM) and sodium pyruvate (1mM) and the pH of the assay media was maintained at 7.4, and prewarmed to 37°C before usage. The test compound 6b had 8X stock concentration in their respective media and was prepared for microplate injections in port A. For mitochondrial stress test, stock solutions of oligomycin (9 µM or 9X), FCCP (2.5 µM or 10X), rotenone+antimycin A (5.5 µM or 11X) were prepared such that their final concentration was 1 µM, 0.25-1uM, and 0.5 µM, respectively in the glucose enriched assay media. For mitochondrial stress test, port A was injected with test compounds at 14.29 minutes, and oligomycin, FCCP, and rotenone+antimycin A were added after 33.8, 53.35, 72.87 minutes respectively. OCR is measured via Seahorse XFe96® analyzer (Agilent). The mitochondrial parameters for MST were calculated utilizing the Wave 2.4.0 software (Agilent).

### In vivo systemic toxicity in CD-1 mice

Five-week-old CD-1 mice (Charles River) were obtained and acclimatized for one week before initial treatment. Mice (n=5) were grouped randomly based on average body weight. Group 1 served as the control group and was administered the vehicle only. Group 2 mice were given niclosamide, while Group 3 mice were administered compound 6b. Vehicle, niclosamide and compound 6b were injected intraperitoneally for a total of 14 days, following a dose-escalation regimen. Mice body weights were recorded daily and euthanized according to IACUC guidelines at the end of the study. A graph of days of treatment versus body weight ± SEM was generated using GraphPad software.

### In vivo efficacy in 67NR syngeneic model in Balb/C mice

5x10^6^ cells 67NR cells suspended in 1:1 matrigel-PBS were injected on back of female Balb/C mice (Charles River). Treatment was initiated when the average tumor volume was ∼100mm^3^. The mice were randomly assigned into groups (n=8) based on average tumor volume and average body weight. Tumors were measured with calipers every two or three days and tumor volumes were calculated using the formula V = (ab^2^)/2 where ‘a’ is the long diameter of the tumor and ‘b’ is the short diameter of the tumor. Mice were euthanized according to IACUC guidelines at the end of the study and tumors were isolated and weighed.

### Ethical Considerations

The experimental procedures involving animals that were conducted at the University of Minnesota Duluth campus, adhering to the U.S. National Institutes of Health Guide for the Care and Use of Laboratory Animals and were approved by the Institutional Animal Care and Use Committee (IACUC). Studies under protocol number 1311-31063A (systemic toxicity **Figure 3A**) and 1605-33796A (67NR syngeneic model, **Figure 3B**) were conducted at the University of Minnesota.

### Statistical Analysis

Statistical analyses were performed using GraphPad Prism version 6.0. Data were presented as average ± SEM of at least three independent experiments. To determine the significance of differences between treated and control groups, one-way ANOVA followed by Dunnett’s multiple comparisons test was conducted. Dose-response curves were fitted using nonlinear regression analysis to calculate the IC50 values. Mann-Whitney test was used to compare the compound-treated and control groups for *in vivo* studies. A P-value of < 0.05 was considered statistically significant, indicating that the observed differences were unlikely due to random chance.

### APPENDIX

#### Spectral Characterization

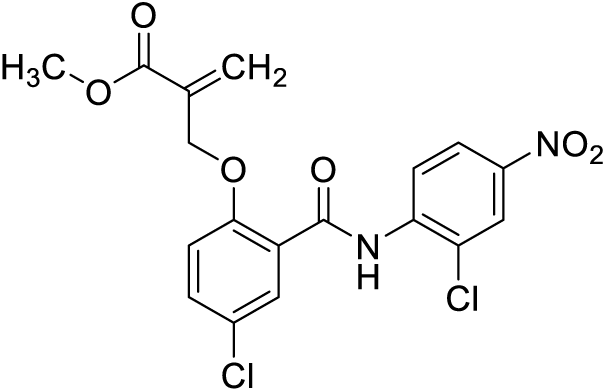

methyl 2-((4-chloro-2-((2-chloro-4-nitrophenyl)carbamoyl)phenoxy)methyl)acrylate

**^1^H NMR (400MHz, DMSO-*d*_6_):** δ 10.57 (s, 1H), 8.66 (d, *J*=9.2 Hz, 1H), 8.43 (d, *J*=2.6 Hz, 1H), 8.29 (dd, *J*=2.64, 9.2 Hz, 1H), 7.98 (d, *J*=2.8 Hz, 1H), 7.68 (dd, *J*=2.84, 8.92 Hz, 1H), 7.4 (d, *J*=9 Hz, 1H), 6.4 (s, 1H), 6.14 (s, 1H), 5.13 (s, 2H), 3.66 (s, 3H)

**^13^C NMR (100MHz, DMSO-*d*_6_):** δ 165.62, 162.67, 155.28, 143.54, 141.15, 135.06, 134.11, 131.06, 130.75, 126.16, 125.22, 124.13, 123.82, 123.36, 122.33, 116.99, 69.00, 52.50

**HRMS (ESI) m/z**: calc’d for C_18_H_14_Cl_2_N_2_O_6_ [M+nH]: 425.0301, found 425.0302

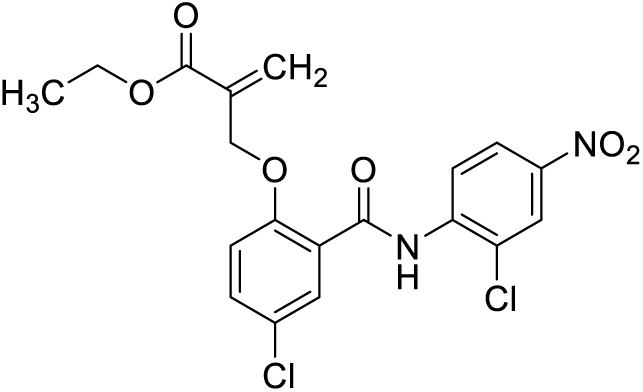

ethyl 2-((4-chloro-2-((2-chloro-4-nitrophenyl)carbamoyl)phenoxy)methyl)acrylate

**^1^H NMR (400MHz, DMSO-*d*_6_):** δ 10.57 (s, 1H), 8.66 (d, *J*=9.2 Hz, 1H), 8.43 (d, *J*=2.64 Hz, 1H), 8.29 (dd, *J*=2.64, 9.16 Hz, 1H), 7.98 (d, *J*=2.8 Hz, 1H), 7.69 (dd, *J*=2.84, 8.92 Hz, 1H), 7.4 (d, *J*=9 Hz, 1H), 6.39 (s, 1H), 6.14 (s, 1H), 5.12 (s, 2H), 4.11 (q, *J*=7.12, 2H), 1.14 (t, *J*=7.08, 3H)

**^13^C NMR (100MHz, DMSO-*d*_6_):** δ 165.14, 162.67, 155.38, 143.54, 141.15, 135.36, 134.16, 131.07, 130.67, 126.16, 125.21, 124.15, 123.80, 123.30, 122.27, 117.02, 69.16, 61.21, 14.35

**HRMS (ESI) m/z**: calc’d for C_19_H_16_Cl_2_N_2_O_6_ [M+nH]: 439.0471, found 439.0458

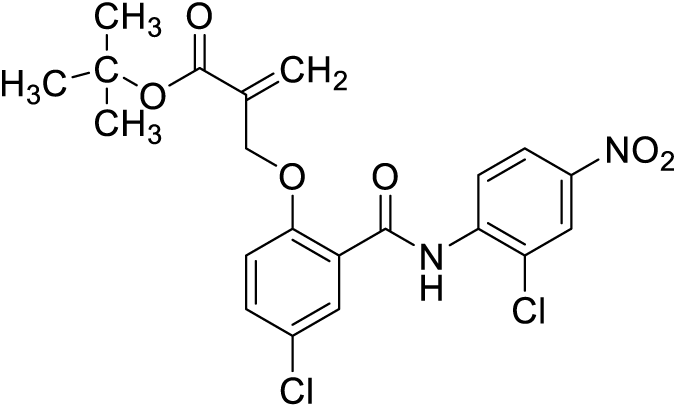

tert-butyl 2-((4-chloro-2-((2-chloro-4-nitrophenyl)carbamoyl)phenoxy)methyl)acrylate

**^1^H NMR (400MHz, CDCl3):** δ 10.57 (s, 1H), 8.68 (d, *J*=9.2 Hz, 1H), 8.44 (d, *J*=2.48 Hz, 1H), 8.31(s, 1H), 8.29 (d, *J*=2.4 Hz, 1H), 7.99 (d, *J*=2.64 Hz, 1H), 7.70 (dd, *J*=2.56, 8.88 Hz, 1H), 7.40 (d, *J*=8.96 Hz, 1H), 6.3 (s, 1H), 6.07 (s, 1H), 5.06 (s, 2H), 1.33 (s, 9H)

**^13^C NMR (100MHz, CDCl3):** δ 164.04, 162.42, 154.90, 143.04, 141.16, 136.02, 133.90, 132.53, 128.15, 127.72, 124.72, 123.56, 122.97, 122.55, 121.17, 114.90, 82.18, 68.95, 28.00

**HRMS (ESI) m/z**: calc’d for C_21_H_20_Cl_2_N_2_O_6_ [M+nH]: 467.0760, found 467.0771

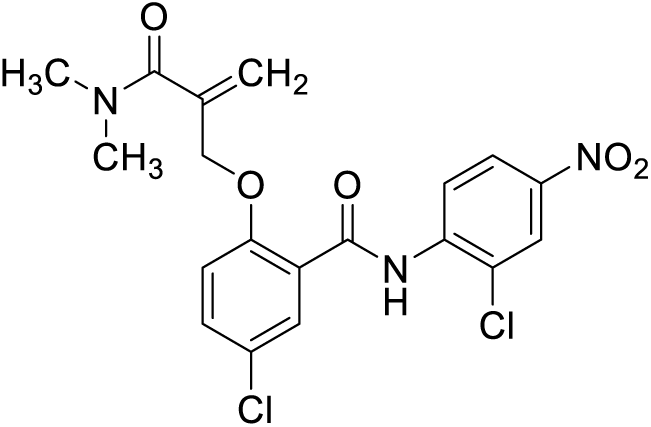

5-chloro-N-(2-chloro-4-nitrophenyl)-2-((2-(dimethylcarbamoyl)allyl)oxy)benzamide

1H **NMR (400MHz, DMSO-*d*_6_):** δ 10.72 (s, 1H), 8.64 (d, *J*=9.16 Hz, 1H), 8.44 (d, *J*=2.64 Hz, 1H), 8.30 (dd, *J*=2.52, 9.08 Hz, 1H), 7.95 (d, *J*=2.8 Hz, 1H), 7.68 (dd, *J*=2.84. 8.96 Hz, 1H), 7.44 (d, *J*=9 Hz, 1H), 5.69 (s, 1H), 5.4 (s, 1H), 5.16 (s, 2H), 2.82 (s, 3H), 2.74 (s, 3H)

**^13^C NMR (100MHz, DMSO-*d*_6_):** δ 168.54, 162.68, 155.13, 143.58, 141.30, 139.08, 133.89, 130.90, 126.16, 125.26, 124.13, 123.79, 122.32, 119.84, 117.21, 70.77, 38.59, 34.70

**HRMS (ESI) m/z**: calc’d for C_19_H_17_Cl_2_N_3_O_5_ [M+nH]: 438.0636, found 438.0618

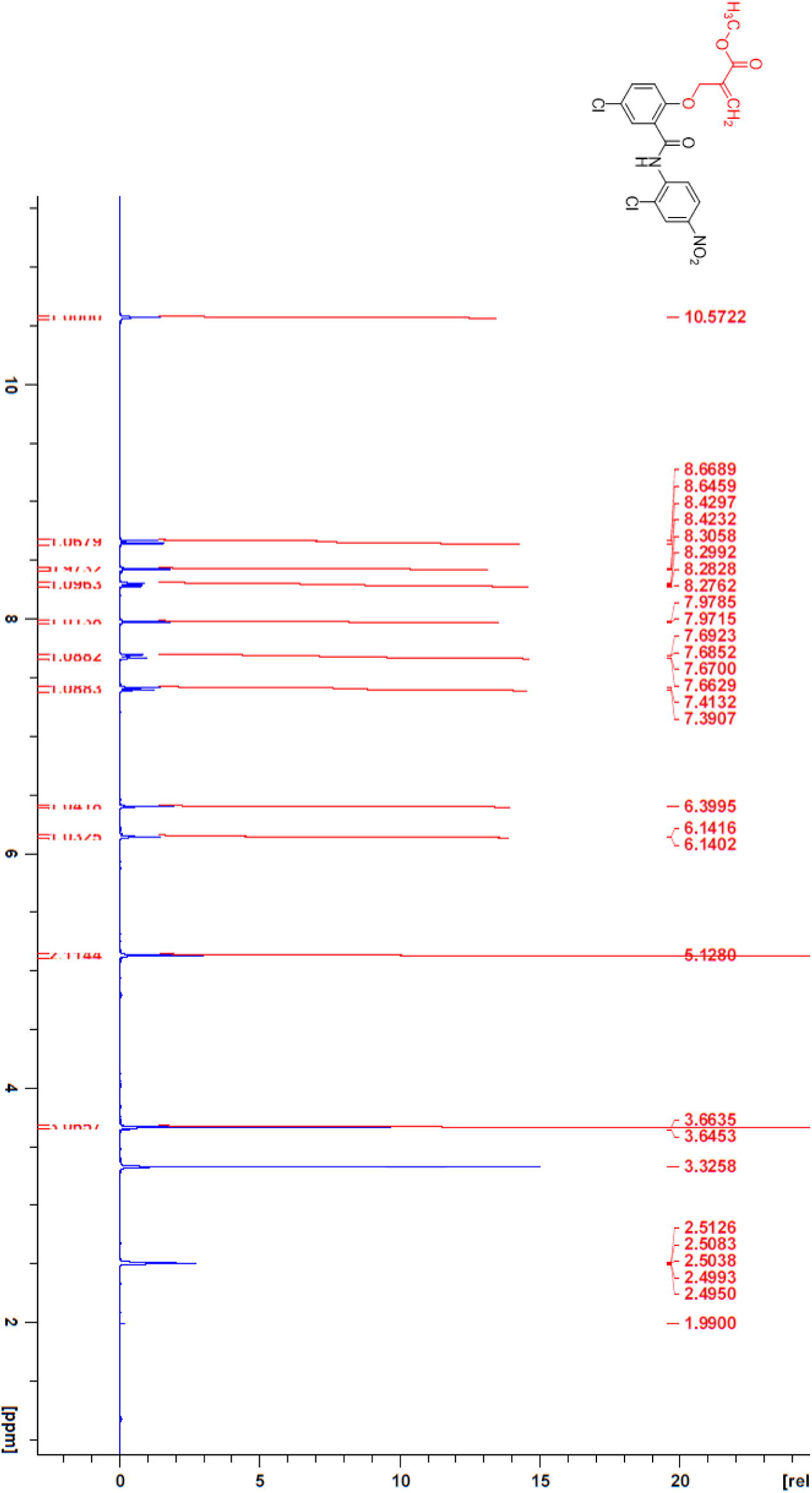

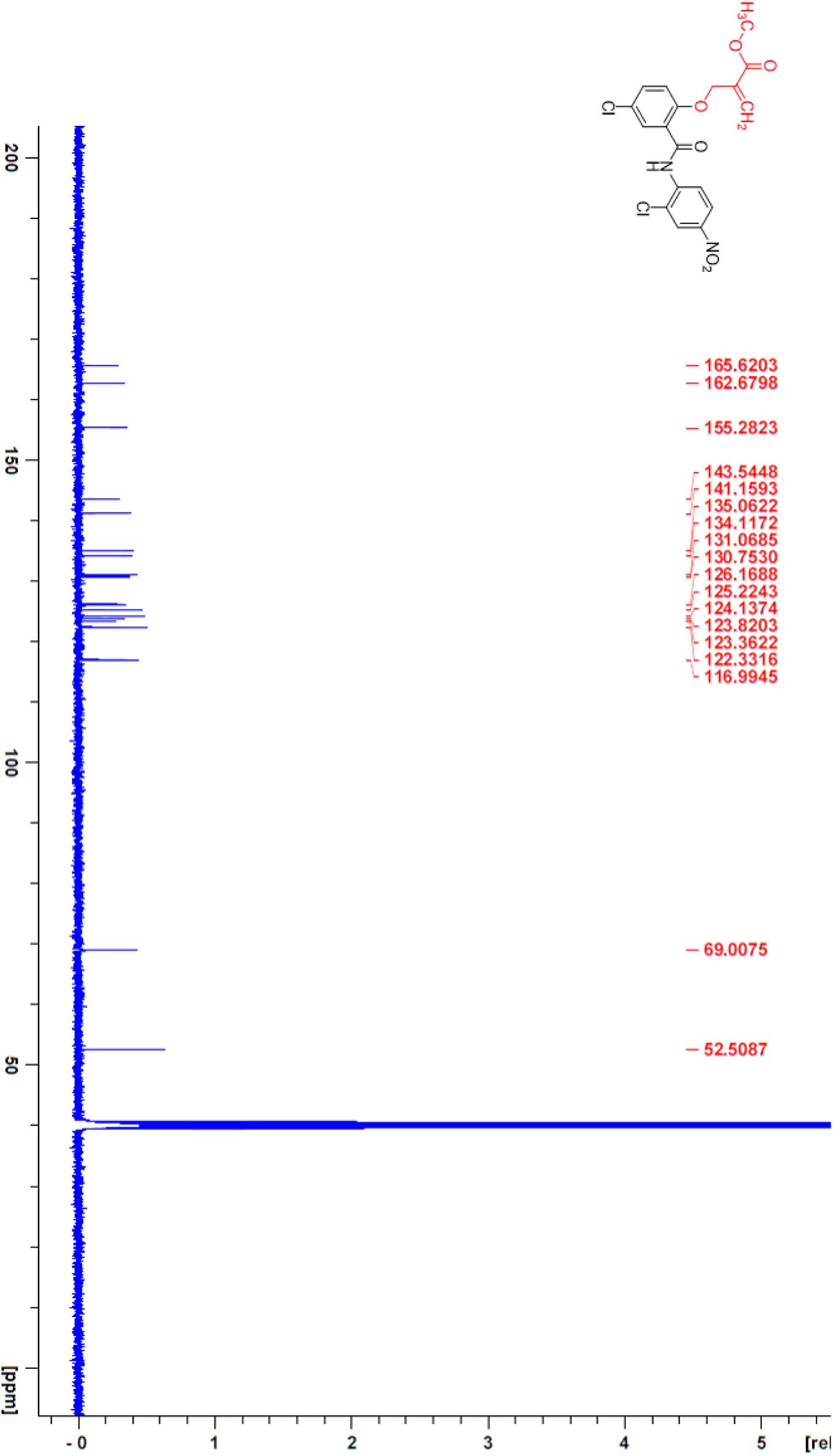

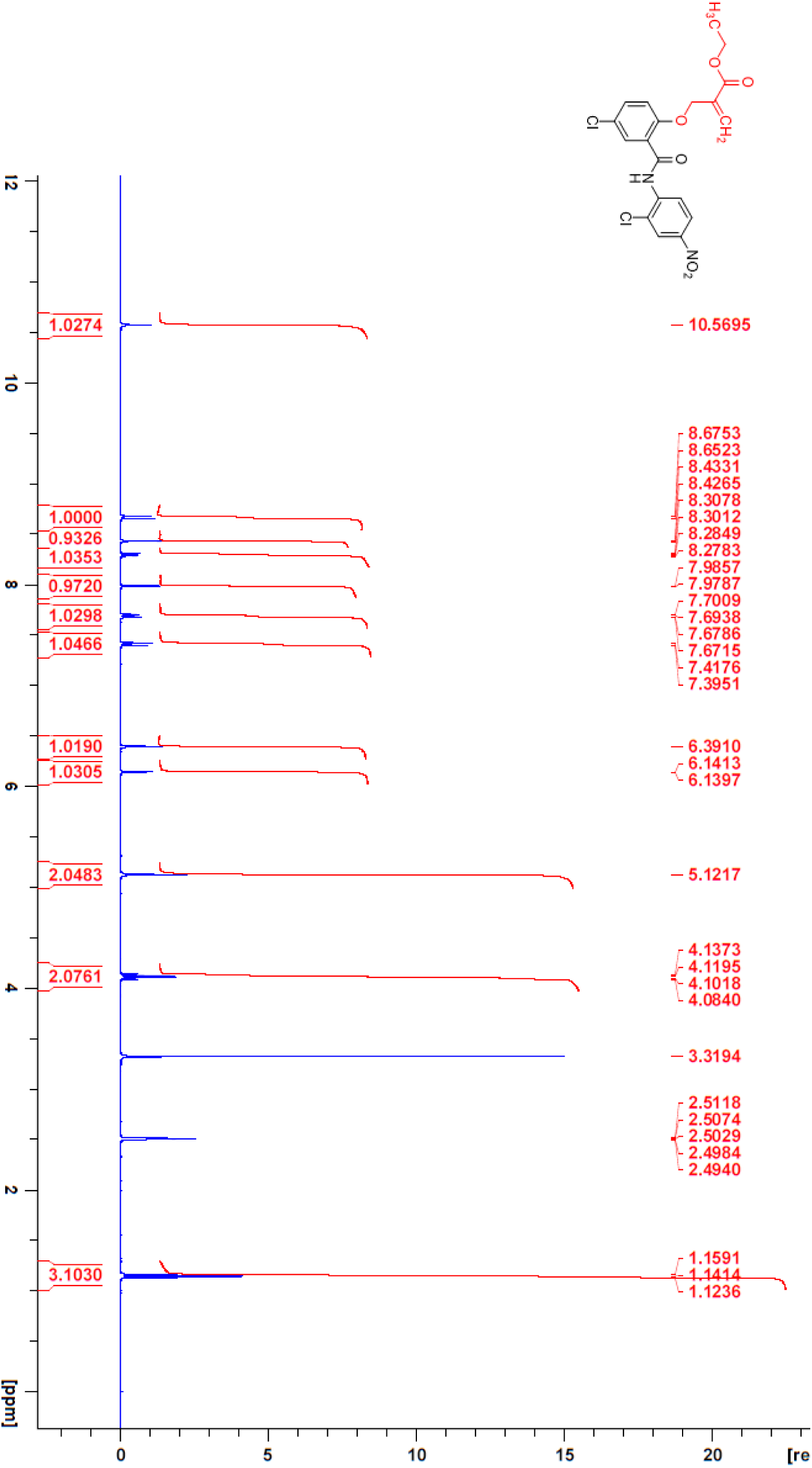

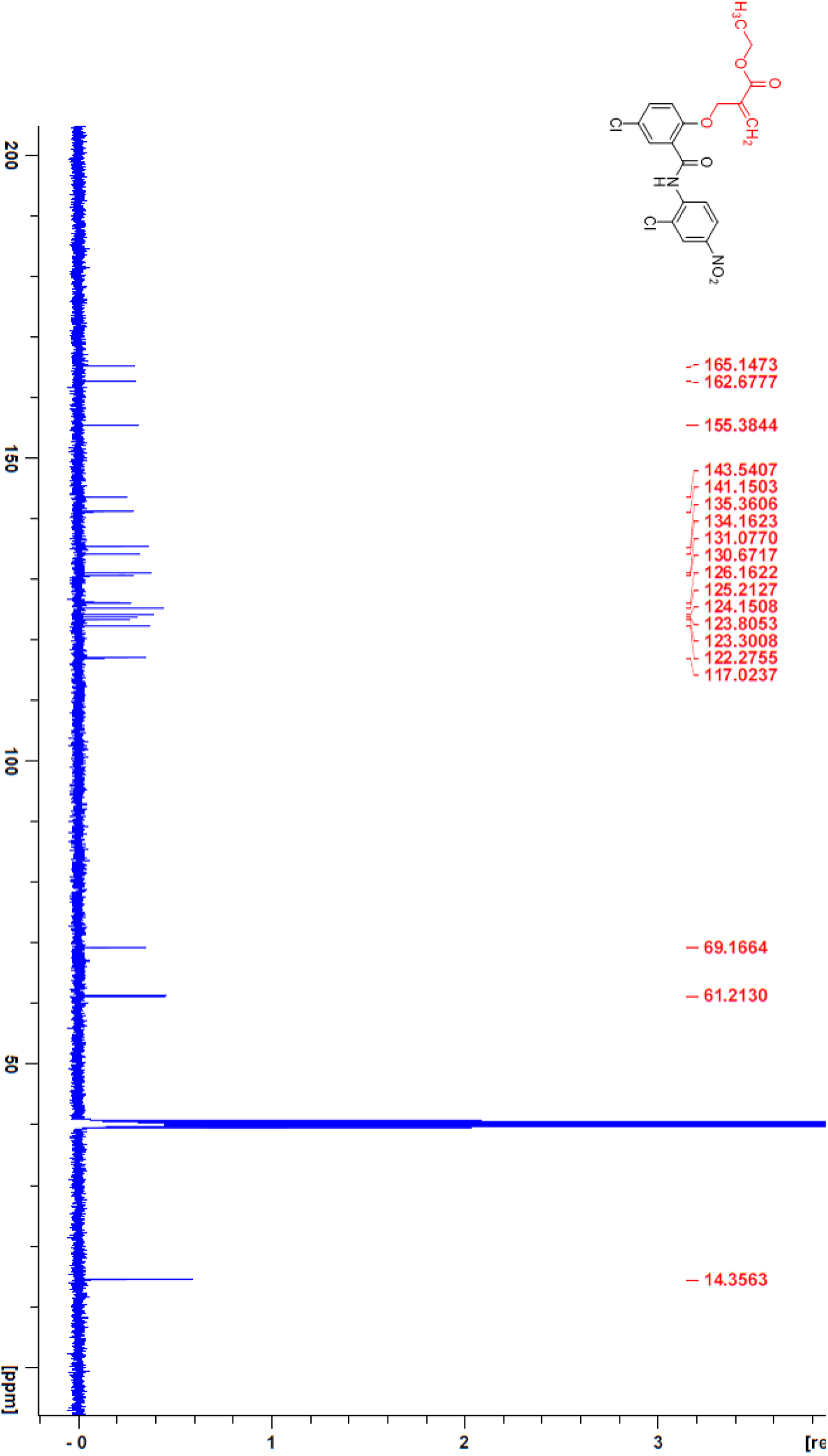

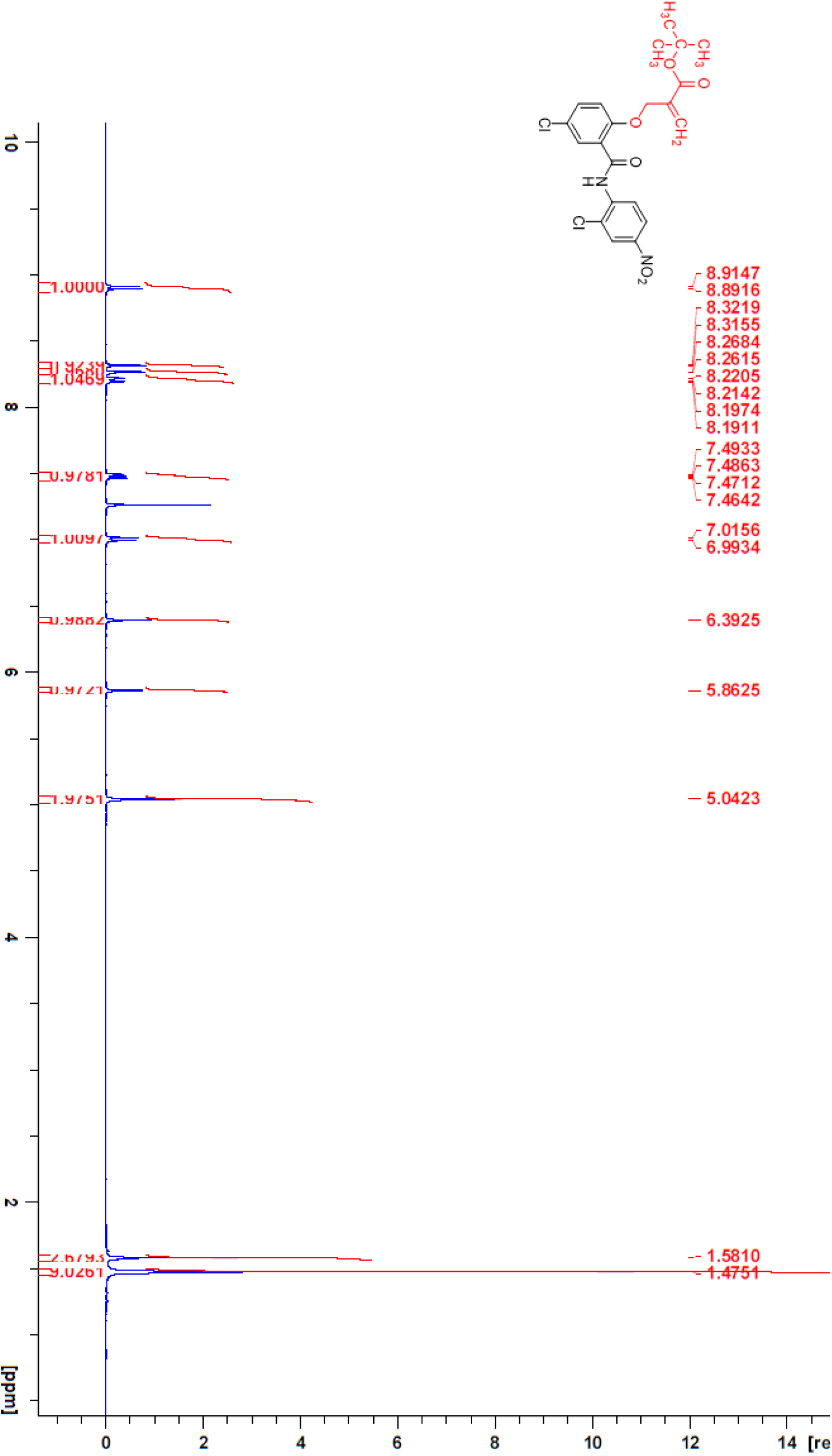

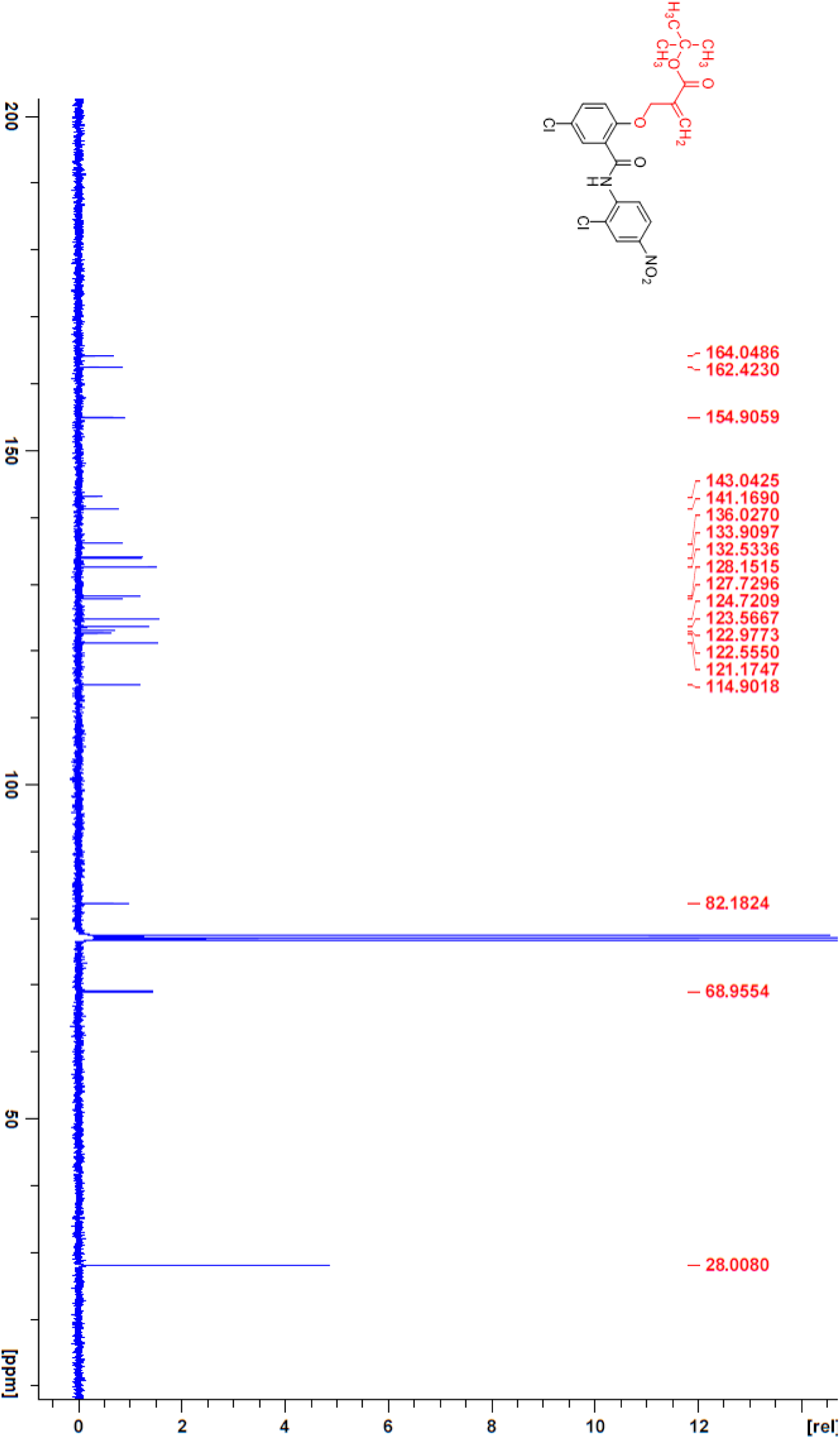

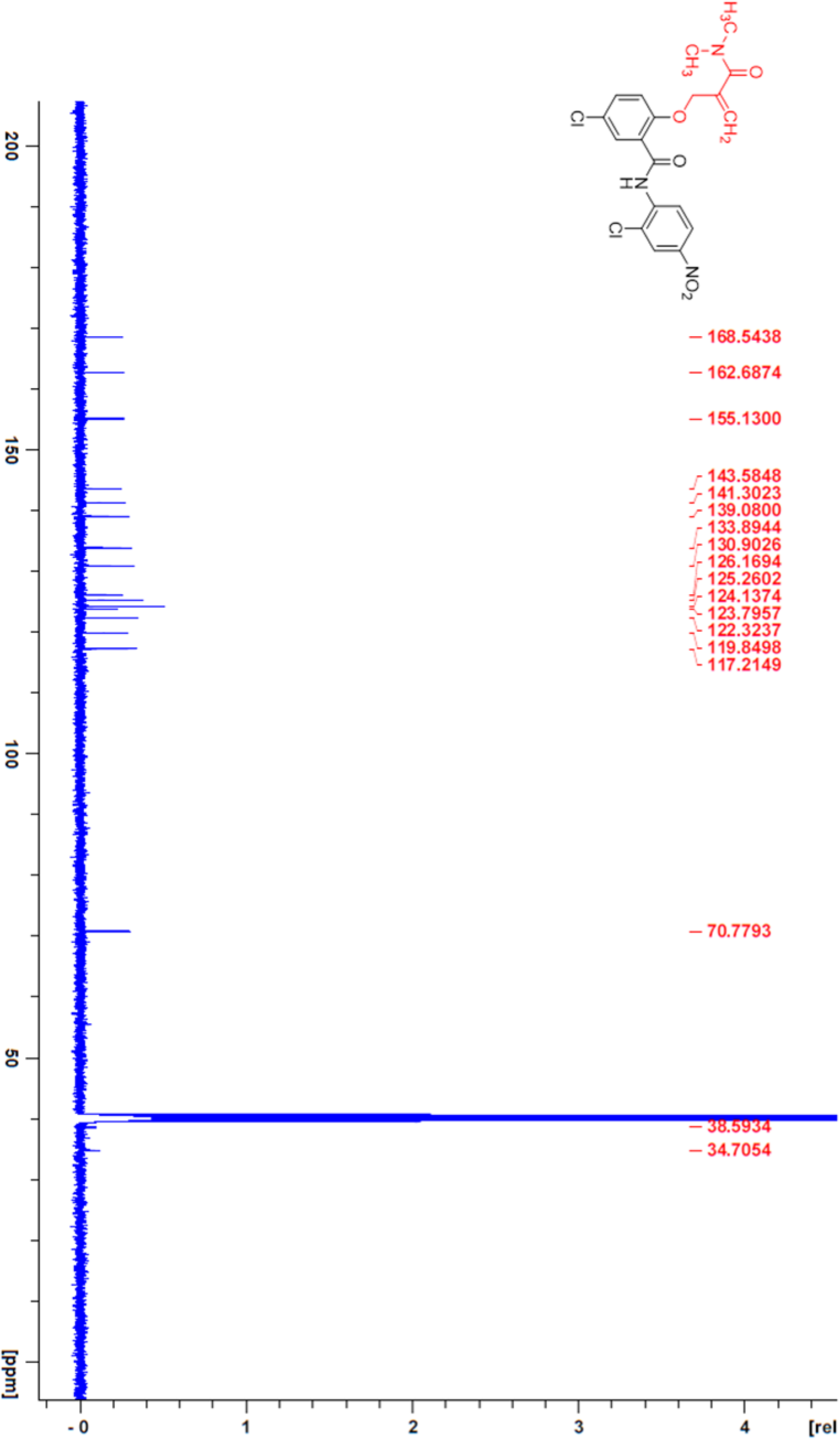

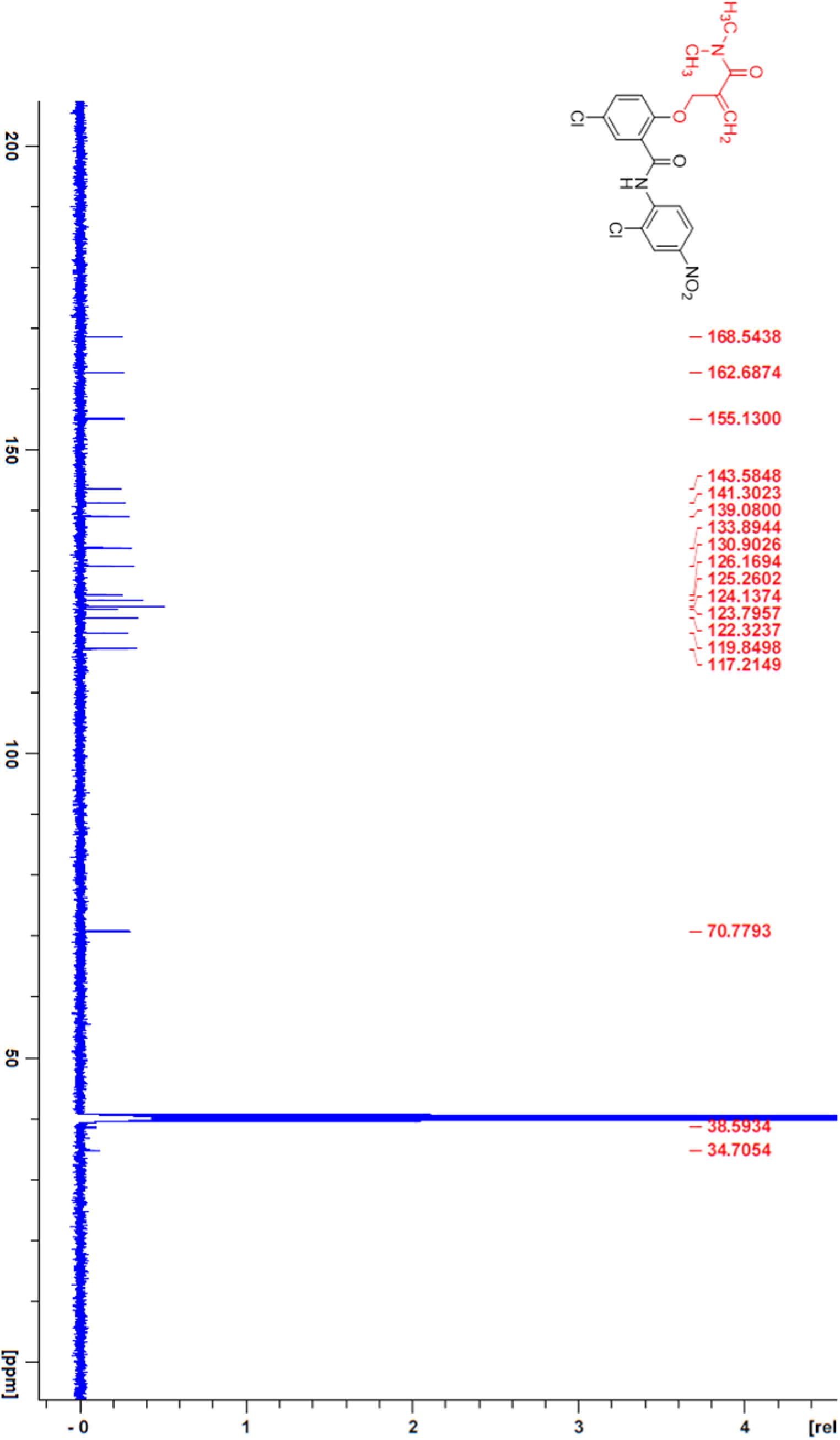

